# Reward signaling in a recurrent circuit of dopaminergic neurons and Kenyon cells

**DOI:** 10.1101/357145

**Authors:** Radostina Lyutova, Maximilian Pfeuffer, Dennis Segebarth, Jens Habenstein, Mareike Selcho, Christian Wegener, Andreas S. Thum, Dennis Pauls

## Abstract

Dopaminergic neurons in the brain of the *Drosophila* larva play a key role in mediating reward information to the mushroom bodies during appetitive olfactory learning and memory. Using optogenetic activation of Kenyon cells we provide evidence that a functional recurrent signaling loop exists between Kenyon cells and dopaminergic neurons of the primary protocerebral anterior (pPAM) cluster. An optogenetic activation of Kenyon cells paired with an odor is sufficient to induce appetitive memory, while a simultaneous impairment of the dopaminergic pPAM neurons abolishes memory expression. Thus, dopaminergic pPAM neurons mediate reward information to the Kenyon cells, but in turn receive feedback from Kenyon cells. We further show that the activation of recurrent signaling routes within mushroom body circuitry increases the persistence of an odor-sugar memory. Our results suggest that sustained activity in a neuronal circuitry is a conserved mechanism in insects and vertebrates to consolidate memories.

## 2. Introduction

Memory can be defined as a change in behavior due to experience. It enables animals and humans to adapt to a variable environment. To do so, the current situation has to be continuously re-evaluated and recorded in the brain. On the neuronal level, memories are encoded as changes in activity or connectivity which outlast the triggering environmental stimulus. Thus, the storage of relevant information in a memory trace is a multi-dimensional and dynamic process. Such complex calculations require a neuronal network with feedforward and feedback motifs enabling the system to integrate new information, to provide learned information, and to organize decision making for the behavioral outcome based on an integration of all information.

In *Drosophila*, the mushroom bodies (MBs) represent a multimodal integration center incorporating a variety of different sensory stimuli. A major function of the MBs is to establish, consolidate and recall associative odor memories, in both the adult and larval *Drosophila*^1–8^. During classical olfactory conditioning, odor information (CS, conditioned stimulus) is represented by an odor-specific subset of third order olfactory neurons, the MB intrinsic Kenyon cells (KCs). KCs also receive information about reward or punishment (US, unconditioned stimulus) mediated by octopaminergic neurons (OANs) and/or dopaminergic neurons (DANs). The coincidental activation of KCs via odor stimulation and MB input neuron (MBIN; OANs or DANs) activation leads to memory formation^1,2,4,5,9,10^.

The larval MB consist of eleven distinct compartments which are defined by the innervation pattern of MBINs and MB output neurons (MBON)^11,12^. The four compartments of the MB medial lobe are each innervated by a single DAN of the pPAM (primary protocerebral anterior medial) cluster. These are essential to mediate the internal reward signal to the medial lobes of the MBs during odor-sugar learning^12–14^. In addition to the four dopaminergic pPAM neurons, OANs are also required for appetitive odor memories in the larva as they mediate the sweetness of a given sugar^15–17^. Yet, it is not clear whether OANs mediate the rewarding sweetness directly to the MBs or onto the dopaminergic pPAM neurons. The finding that OANs do not innervate the medial lobes of the MBs in the larva, but potentially pPAM neurons, suggests that OANs might function directly upstream of the dopaminergic pPAM neurons as suggested in adult *Drosophila*^13,16–19^.

The current assumption is that the MBs are the brain region where synaptic plasticity occurs and from where plasticity is transferred into a conditional behavioral output driven by MBONs and downstream premotor circuits^2,4,5,10^. Recently, Eichler and coworkers reconstructed the connectome of the larval MBs providing a complete neuronal and even synaptic circuit map of KCs, MBINs, and MBONs on the electron-microscopic level^11^. Interestingly, this study describes a canonical circuit motif with coherent characteristics in each compartment of the MBs. This motif consists of known connections – MBINs with synaptic connections to KCs and synaptic connections of KCs onto MBONs; but it also revealed new, so far unexpected contacts: (a) reciprocal synaptic connections between KCs; (b) recurrent synaptic connections from KCs onto MBINs, and (c) direct connections from MBINs onto MBONs. So far, the function of these new connections is largely unexplored, especially in the larva.

It is, however, tempting to speculate that feedback signaling within the MB circuitry allows modulation of neuronal activity on different levels. This is supported by recent findings in adult *Drosophila,* where feedback signaling to MBINs is important for appetitive olfactory long-term memory formation^20,21^.

Here, we experimentally address whether recently anatomically described larval KC-to-DAN synapses are functional and test whether they have a role in appetitive olfactory learning in the *Drosophila* larva. We find that optogenetic activation of MB KCs paired with odor stimulation induces an appetitive memory, which is dependent on KC-to-DAN signaling. Moreover, calcium imaging shows that dopaminergic pPAM neurons specifically respond with an increase in Ca^2+^ levels to MB KC activation. Our results indicate that a persistent recurrent activity within the neuronal network of the MBs helps to stabilize appetitive odor memories in the *Drosophila* larva.

## 3. Results

We used a well-established 1-odor reciprocal training regime to analyze classical olfactory learning and memory in *Drosophila* larvae 9,22. To address whether optogenetic activation of MB KCs affects learning and memory, we used the Gal4/UAS system^23^ to express Channelrhodopsin (UAS-*ChR2-XXL;*^24^) using the *H24-Gal4* driver line ^25^. *H24-Gal4* specifically expresses Gal4 in all KCs and only a few neurons located in the ventral nervous system (VNS) (Fig.1A^II^).

In contrast to the standard procedure, olfactory stimuli were paired with a blue light-dependent activation of KCs (KC-substitution learning) (Fig. 1A, 1A^I^) as an artificial substitute for a conventional US like sugar or salt^5,9^. Strikingly, experimental larvae showed a significant appetitive memory in contrast to genetic controls that did not show any learning performance (Fig. 1A^III^, Fig. S1). This result suggests that the artificial activation of KCs is sufficient to induce an appetitive memory. To validate our findings and to exclude that the effect is based on the activation of cells of the VNS in *H24>ChR2-XXL* larvae, we additionally tested *OK107-Gal4* (Fig. 1B) in the KC-substitution learning experiment. Similar to *H24-Gal4*, *OK107-Gal4* was shown to label all KCs^26^. Again, artificial activation of KCs using *OK107-Gal4* led to an appetitive memory formation in experimental larvae (Fig. 1B^I^). Thus, KC-substitution learning using two MB-specific Gal4 lines confirmed that optogenetic activation of KCs is sufficient to induce an appetitive memory (Fig. 1A, 1B).

**Fig. 1:**
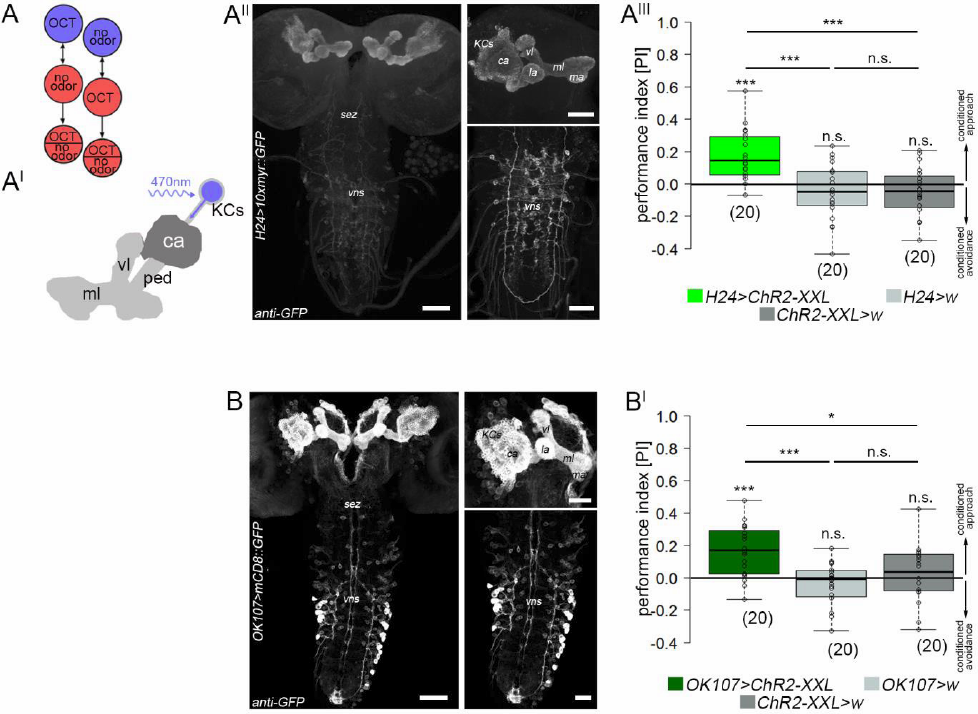
Optogenetic activation of mushroom body Kenyon cells during conditioning is sufficient to induce appetitive memory formation. (A) Illustration of the 1-odor reciprocal training regime used in this study (**substitution learning**). (A^I^) Schematic drawing of the larval MB expressing ChR2-XXL in all KCs. (A^II^) Expression pattern of H24-Gal4 crossed with 10xUAS-myr::GFP stained with anti-GFP (white). H24-Gal4 shows expression in the complete MBs and a few additional cells of the VNS. (A^III^) Conditional optogenetic activation of KCs coupled to odor information during training is sufficient to induce an appetitive memory. (B) Expression pattern of OK107-Gal4 crossed with UAS-mCD8::GFP stained with anti-GFP (white). (B^I^) Conditional optogenetic activation of KCs coupled to odor stimulation during training is sufficient to induce an appetitive memory expression in OK107-Gal4. ca: calyx; ChR2-XXL: channelrhodopsin2-XXL; GFP: green-fluorescent protein; KCs: Kenyon cells; la: lateral appendix; ma: medial appendix; ml: medial lobe; OCT: octanol; ped: peduncle; sez: subesophageal zone; vl: vertical lobe; vns: ventral nervous system; w: w^1118^. Significance p>levels: p>0.05 n.s., p<0.05 *, p<0.01 **, p<0.001 ***. Scale bars: 50 μm and 25 μm for higher magnifications.

### Optogenetic activation of mushroom body Kenyon cells does not affect larval locomotion or innate odor responses

The MBs integrate a variety of different sensory modalities. To exclude that the optogenetic activation of KCs alters general behaviors like locomotion or the processing of sensory stimuli necessary for classical olfactory learning (i.e. odors), larvae were assayed for (i) locomotion and (ii) innate odor preference. We analyzed larval locomotion using the FIM tracking system^27^. For this, larvae were monitored under red light for one minute. Subsequently, larvae were exposed to blue light and recorded again for one minute (Fig. 2; the light regime is indicated by the red and blue rectangles). Optogenetic activation of KCs did not affect the velocity and crawled distance over time of *H24>ChR2-XXL* larvae, suggesting that artificial activation of KCs does not affect general locomotion parameters (Fig. 2A-A^II^, Fig. S2). However, experimental larvae showed a significant decrease in the number of turns and stops under blue light exposure, suggesting a change in orientation behavior (Fig. S2^I^, S2^II^). Next, we tested whether the optogenetic activation of KCs alters innate odor preference, which would perturb the conditioning experiments of our study. For this, larvae were assayed in a simple choice test using pure octanol (OCT; the odor used in our KC-substitution learning) and diluted amylacetate (AM; commonly used for the two-odor reciprocal experimental design;^5,9^), respectively. Optogenetic activation of KCs in experimental larvae did not change the innate preference to OCT and AM (Fig. 2B, 2B^I^). To verify that the activation of all KCs does not change odor quality to an unspecific attractive odor, larvae were tested in a choice test with exposure to OCT and AM opposing each other. Previous studies showed that pure OCT and 1:40 diluted AM are balanced in their attractiveness to larvae, while larvae prefer pure AM over pure OCT. As expected, experimental larvae showed a significant approach towards AM, which was indistinguishable from controls (Fig. 2B^II^); coherently both experimental and control larvae were randomly distributed in the choice test between OCT and 1:40 diluted AM (Fig. 2B^III^). These results suggest that an optogenetic activation of all KCs does not change innate odor preferences. We also repeated the KC-substitution learning experiment using the two-odor reciprocal design (Fig. 2C;^5,9^). Similar to our initial results, optogenetic activation of all KCs was sufficient to induce appetitive memory, verifying that odor discrimination is not disturbed due to artificial activation of KCs via ChR2-XXL (Fig. 2C^I^). Taken together, these results suggest that optogenetic activation of KCs does not perturb locomotion and innate preference to odors essential for KC-substitution learning.

**Fig. 2:**
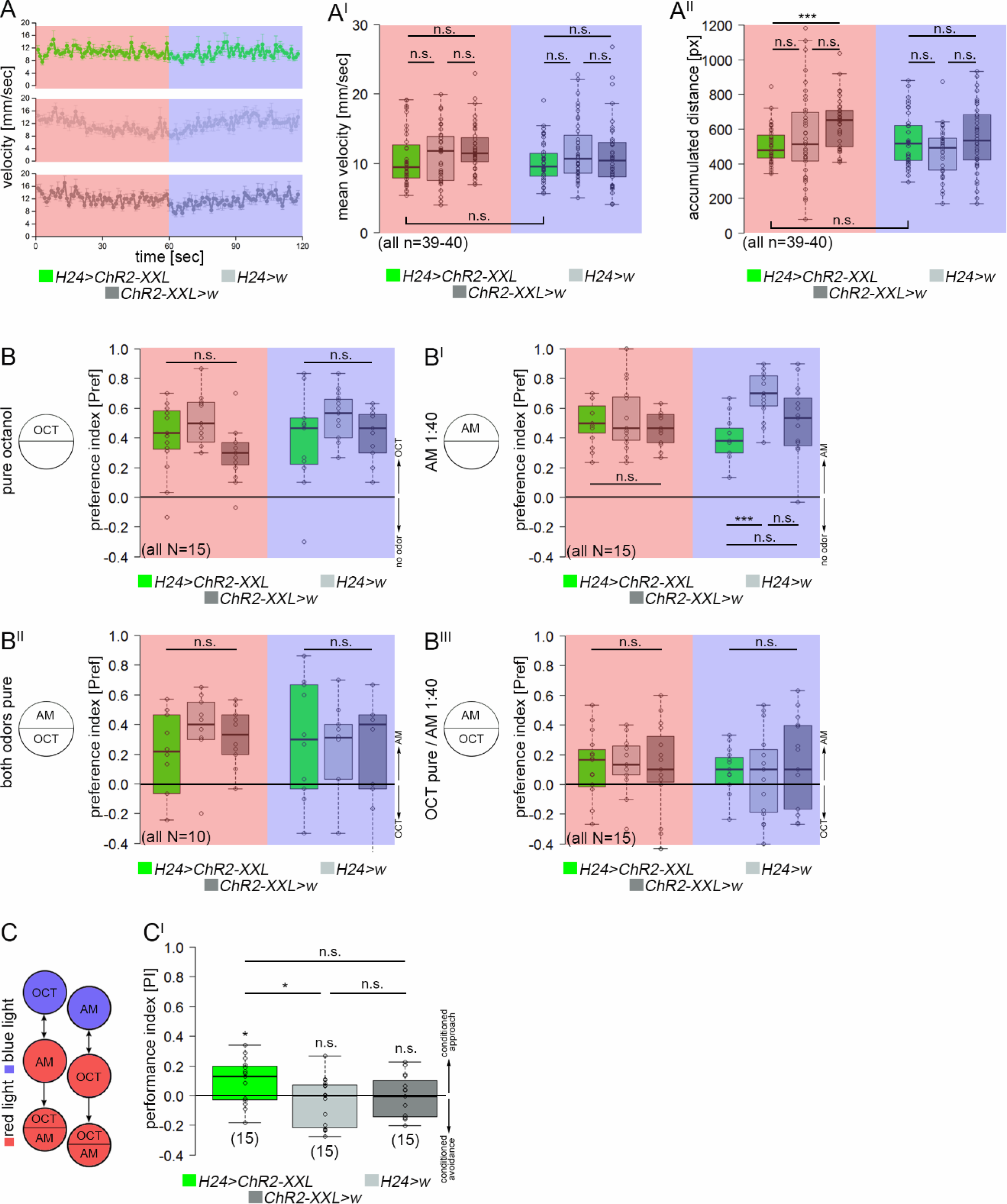
Optogenetic activation of mushroom body Kenyon cells does not alter locomotor behavior or innate odor preference in larvae. (A) Conditional optogenetic activation of KCs does not change larval velocity (A and A^I^) and accumulated distance (A^II^). The light regime is indicated by the red and blue rectangles: larvae were monitored under red light for one minute, subsequently under blue light for another minute. (B) Similarly, innate odor preference to OCT (B) and diluted AM (B^I^) is not altered due to conditional optogenetic activation of KCs. In line, experimental larvae showed normal performance in an odor discrimination task using OCT and either pure AM (B^II^) or 1:40 diluted AM (B^III^). (C and C^I^) Conditional optogenetic activation of KCs is sufficient to induce an appetitive memory using a 2-odor reciprocal training regime indicating that experimental larvae can distinguish different odors despite artificially activated KCs. AM: amylacetate; ChR2-XXL: channelrhodopsin2-XXL; KCs: Kenyon cells; OCT: octanol; w: w^1118^. Significance levels: p>0.05 n.s., p<0.05 *, p<0.01 **, p<0.001 ***.

### Why is the activation of KCs sufficient to specifically induce appetitive memory?

Our results so far illustrate that artificial activation of KCs paired with odor exposure is sufficient to specifically induce appetitive memory. Therefore, we tested whether the optogenetic activation of KCs indeed mimics an internal reward signal. Such an internal reward signal would be sufficient to induce an associative reward memory in our KC-substitution learning experiment if tested on a test plate without US information (Fig. 1A). We tested larvae in a simple light preference test. Feeding larvae strongly avoid light to ensure staying within the food^28–30^. Thus, we challenged *H24>ChR2-XXL* larvae to choose between blue light and darkness. Given the idea that the activation of KCs induces internal reward signaling, we assumed that control larvae would prefer darkness over blue light, whereas *H24>ChR2-XXL* larvae would show a reduced dark preference as they would experience blue light as reward (Fig. 3A). As expected, control larvae significantly avoided the illuminated rectangles after 1min and 3min. In contrast, experimental larvae showed light avoidance only after 3min, but were randomly distributed after 1min suggesting that the experimental larvae showed a delayed light avoidance (Fig. 3A^I^). To verify that larvae are challenged to decide between reward and innate light avoidance in our assay, we repeated the experiment, but this time used red light, which does not elicit innate light avoidance in the larva^31,32^(Fig. 3A^II^), but activates KCs in experimental larvae as we expressed *UAS-Chrimson*, a red-light driven channelrhodopsin^33^. *H24>Chrimson* larvae showed a significant preference for the red-light illuminated rectangles within the first minutes, while control larvae were randomly distributed. Similar to our previous results, the effect was gone after 3min (Fig. 3A^III^). To test this even further, we challenged the larvae to decide between OCT in the darkness and blue light at the opposing side (Fig. S4, S4^I^). In this assay, control larvae are driven to go into the darkness due to their innate light avoidance and innate OCT preference. Control larvae strongly preferred the dark and OCT side over blue light. Experimental larvae, however, showed significantly reduced preference for darkness and OCT. This supports our hypothesis that experimental larvae were challenged to decide between their innate OCT and darkness preference versus an artificially induced internal reward signaling in the blue light (Fig. S4^I^). Taken together, our results suggest that optogenetic activation of KCs elicits an internal reward signaling which induces appetitive memory during KC-substitution learning. This is further supported by the change in orientation behavior, reflected by the reduction in turning rate and direction (Fig. S2I, S2^II^), in larvae with optogenetically activated KCs. Blue light exposure without any gradient may provide an even distributed pleasant environment in experimental larvae due to the optogenetic activation of internal reward signaling. Consequently, experimental larvae reduce turning rate and turning direction, but not the speed (Fig. S2^I^, S2^II^)^34^.

**Fig. 3:**
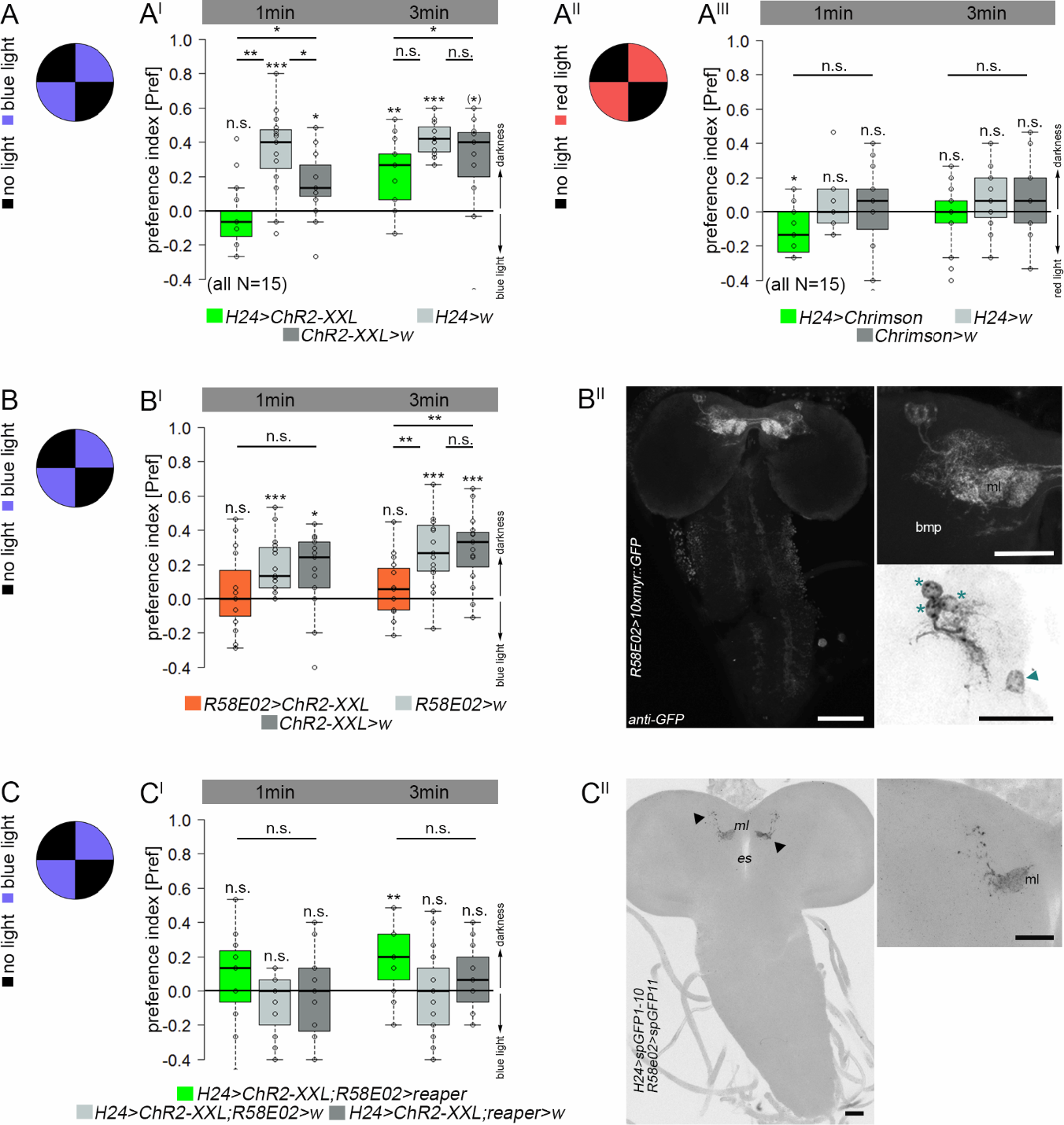
Optogenetic activation of mushroom body Kenyon cells induces internal reward signaling via dopaminergic neurons of the pPAM cluster. Control larvae showed blue light avoidance in a simple choice test (A). H24>ChR2-XXL larvae do not show any light avoidance after 1min, but after 3min (A^I^) suggesting that the blue light-dependent activation of ChR2-XXL induces internal reward signaling. (A^II^ and A^III^) In line, H24>Chrimson larvae prefer red light over darkness after 1min, while genetic controls were distributed randomly. (B+B^I^) Optogenetic activation of pPAM neurons using R58E02-Gal4 abolished light avoidance. (B^II^) Expression pattern of R58E02-Gal4 crossed with 10xUAS-myr::GFP and stained with anti-GFP (white). R58E02-Gal4 shows specific expression in three DANs of the pPAM (protocerebral anterior medial) cluster (asterisks) and one additional cell body at the midline (arrow head). pPAM neurons innervate the medial lobe of the MBs. (C and C^I^) Optogenetic activation of KCs abolished light avoidance in both genetic controls; in contrast experimental larvae with ablated pPAM neurons showed light avoidance after 3min, indicating that optogenetic activation of KCs induces internal reward signaling via pPAM neurons. (C^II^) Reconstituted split-GFP (with anti-GFP) between H24-Gal4 positive neurons and R58E02-Gal4 positive neurons is only visible at the level of the mushroom bodies (arrowheads). bmp: basomedial protocerebrum; ChAT: cholinacetyltransferase; ChR2-XXL: channelrhodopsin2-XXL; DANs: dopaminergic neurons; FasII: fasciclinII; KCs: Kenyon cells; MBs: mushroom bodies; ml: medial lobe; es: esophagus; w: w^1118^. Significance levels: p>0.05 n.s., p<0.05 *, p<0.01 **, p<0.001 ***. Scale bars: 50 μm and 25 μm for higher magnifications.

In our experiments larvae are tested on pure agarose, thus they will just expect a significant gain through the expression of appetitive memory. We argue that the lack of aversive olfactory memory expression is based on the test situation in our KC-substitution learning experiment, as we test larvae intentionally on pure agarose (Fig. 1A). *Drosophila* larvae are known to express aversive memories only in the presence of the aversive US in the test situation^5,9,35^. In contrast, to recall appetitive memories the appetitive US must be absent during the test (Fig. S3). Larvae compare the value of a given US in the test situation with the one used during training and only an expected gain will drive the conditional output behavior^5,9,36^.

### Dopaminergic neurons of the pPAM cluster mediate internal reward signals

Previous studies have shown that dopaminergic cells of the pPAM cluster in the larval and adult *Drosophila* mediate internal reward signals during associative conditioning^12,13,16,37^. To test whether the internal reward signaling described in our study is based on the activity of pPAM neurons, we repeated the light avoidance experiment, but this time expressed ChR2-XXL via *R58E02-Gal4* which labels 3 out of 4 neurons of the larval pPAM cluster (Fig. 3B^II^)^13^. In line with our previous results, optogenetic activation of pPAM neurons abolished light avoidance in *R58E02>ChR2-XXL* larvae (Fig. 3B^I^). The effect was even stronger compared to *H24>ChR2-XXL* larvae (Fig. 3A^I^), as *R58E02>ChR2-XXL* larvae showed no light avoidance after both 1min and 3min (Fig. 3B^I^). Further, we tested whether the pPAM-dependent reward signaling is induced by the optogenetic activation of KCs in *H24>ChR2-XXL* larvae. We used the lexA/LexAop system^38^ together with the Gal4/UAS system to activate KCs (*H24>ChR2-XXL*) and simultaneously ablate pPAM neurons (*R58E02>reaper*). As expected, optogenetic activation of KCs abolished the innate light avoidance in both genetic controls (*H24>ChR2-XXL,R58E02>w* and *H24>ChR2-XXL;reaper>w*) after 1min and 3min (Fig. 3C^I^). In contrast, experimental larvae with ablated pPAM neurons showed light avoidance after 3min, indicating that optogenetic activation of KCs induces internal reward signaling via the activation of pPAM neurons. These results are supported by the electron-microscopic reconstruction of the larval MBs, which described a canonical circuit motif in each compartment including direct KC-to-MBIN synaptic connections^11^. In line, we used the GRASP technique which allows to reconstitute potential synaptic connections as GFP fluorescence is only visible when the neurons of interest are in close vicinity^39^. GFP fluorescence was exclusively detectable in the MBs, mostly in the medial lobe which is the main output region of the pPAM neurons (Fig. 3C^II^)^13^. This supports the hypothesis that the optogenetic activation of KCs induces pPAM-dependent reward signaling directly at the level of the MBs, as no signal was elsewhere detectable.

### Kenyon cell to dopaminergic neuron signaling is sufficient to establish an appetitive memory

Our results suggest that the optogenetic activation of KCs induces reward signaling involving a recurrent pPAM cluster loop. Previous studies already showed that the DANs of the pPAM cluster are important MBINs to establish larval olfactory memories in the *Drosophila* larva^12,13^. Thus, we tested whether the recurrent KC-to-pPAM loop is important to establish the appetitive memory observed in our KC-substitution learning experiment (Fig. 1A^III^). First, we tested whether the ablation of pPAM neurons affects appetitive memory established via optogenetic activation of KCs (*H24>ChR2-XXL;R58E02>reaper*). Indeed, experimental larvae showed no appetitive memory in contrast to the genetic controls (Fig. 4A^I^). Second, to verify these results, we activated the KCs and simultaneously downregulated the dopamine receptor DopR1 (*dcr*;*H24>ChR2-XXL;DopR1-RNAi*) in all KCs. DopR1 is necessary for appetitive olfactory learning^14,40^. Experimental larvae showed no appetitive memory in contrast to the genetic control (*dcr;H24>ChR2-XXL*; Fig. 4B^I^). Importantly, impaired DA signaling did not affect learning-relevant odor preferences (Fig. 4A^II^, 4B^II^). Next, we monitored changes of intracellular Ca^2+^ levels in R58E02-positive pPAM neurons upon optogenetic activation of KCs (*H24>ChR2-XXL;R58E02>GCamp6m*). Isolated brains were first exposed with a low blue light intensity (˜700μW/cm^2^) to monitor baseline activity; subsequently, brains were exposed with a higher light intensity (˜3800μW/cm^2^). This light intensity was sufficient to induce a significant increase in Ca^2+^ levels within pPAM neurons due to the activation of ChR2-XXL in KCs, which confirms the functionality of the KC-to-PAM connection (Fig. 4C: exemplary responses; 4C^I^). Coherently, we found a similar increase in intracellular Ca^2+^ levels in pPAM neurons due to the optogenetic activation of KCs by using UAS-*Chrimson* (Fig. S5, S5^I^). Taken together, our results suggest that functional recurrent signaling between MB KCs and pPAM neurons exist in the *Drosophila* larva.

**Fig. 4:**
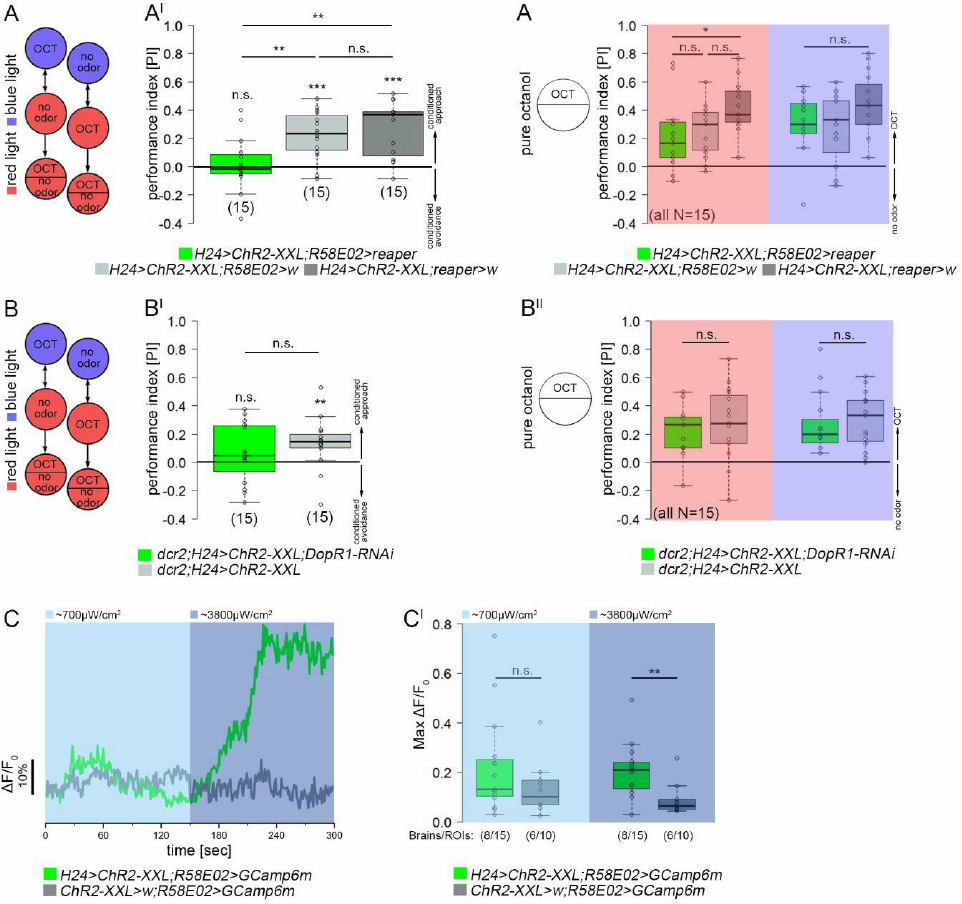
Appetitive memory expression due to the optogenetic activation is based on Kenyon cells to dopaminergic pPAM neurons signaling. (A and A^I^) Optogenetic activation of KCs and simultaneous ablation of pPAM neurons via the expression of the apoptotic gene reaper impairs optogenetically induced appetitive memory. (A^II^) Innate odor preference is unchanged in experimental larvae under red and blue light. (B and B^I^) Similarly, optogenetic activation of KCs and simultaneous knockdown of the dopamine receptor DopR1 impairs appetitive memory formation. (B^II^) Innate odor preference is unchanged in experimental larvae under red light (red rectangle) and blue light (blue rectangle). (C and C^I^) pPAM neurons respond with a significant increase in intracellular Ca^2+^ levels (C^I^) to an optogenetic activation of MB KCs (green; illumination with blue light (475nm)) verifying a functional KC-to-DAN feedback connection. An increase in Ca^2+^ levels was absent in genetic controls (grey) indicating that the response in the experimental brains was specific to the optogenetic activation rather than to light per se. ChR2-XXL: channelrhodopsin2-XXL; KCs: Kenyon cells; MBs: mushroom bodies; OCT: octanol; w: w^1118^. Significance levels: p>0.05 n.s., p<0.05 *, p<0.01 **, p<0.001 ***.

### Reward signaling in the recurrent MB circuit stabilizes appetitive memories over time

The larval MB connectome revealed a complex canonical circuit motif in each MB compartment, with suggested MBIN-to-KC and KC-to-MBON synapses, but also unknown recurrent KC-to-MBIN and MBIN-to-MBON synapses^11^. Our data suggests that the KC-to-MBIN connection is functional, which is in line with a recent report in adult *Drosophila*^41^. To test whether KC-to-pPAM feedback signaling affects normal appetitive olfactory learning, we exposed experimental larvae to a real sugar stimulus and a simultaneous optogenetic activation of KCs via ChR2-XXL, while control flies were trained with sugar only under red light without any blue light exposure (Fig. 5A). *H24>ChR2-XXL* larvae trained to associate OCT with sugar under red light (normal sugar learning) showed significant appetitive memory expression immediately after training. However, memory expression was abolished 15min after training (Fig. 5A^II^, 5B). *H24>ChR2-XXL* larvae that received sugar stimulation in combination with artificial KC activation during training showed comparable appetitive memory expression immediately after training. However, the appetitive memory was still detectable 45min after training, suggesting a change in memory persistence based on artificial activation of the recurrent MB network (Fig. 5A^II^, 5B). Optogenetic activation of KCs did not change innate preference for sugar in *H24>ChR2-XXL* larvae (Fig. S6, S6^I^).

**Fig. 5:**
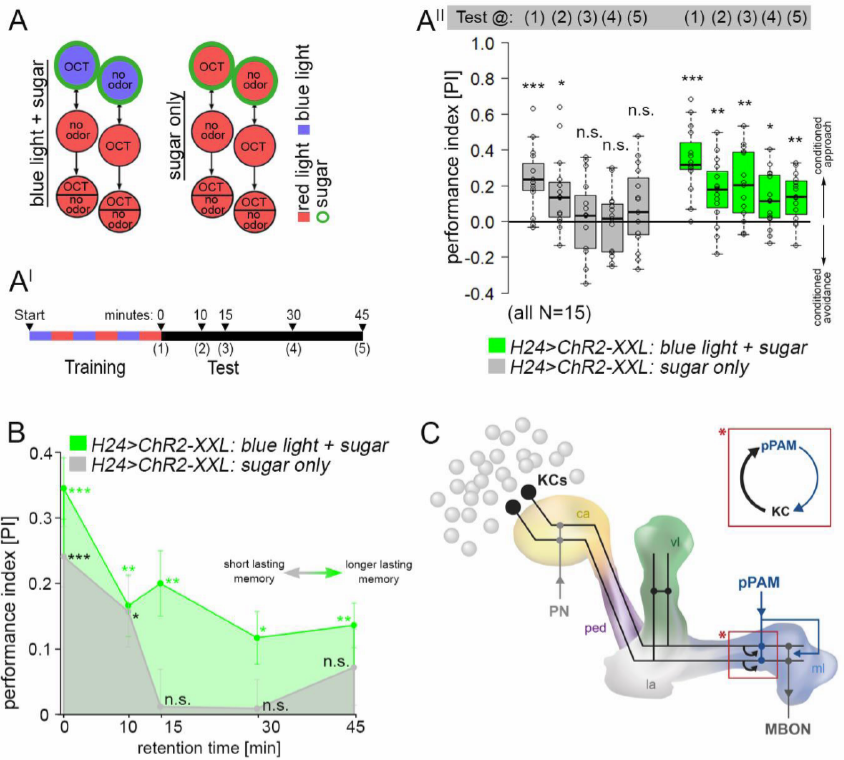
Sustained activity in the mushroom body circuit increases memory stabilization. (A and A^I^) Schematic drawing of the training and test regime: H24>ChR2-XXL larvae were either trained for odor-sugar associations under red light, or for odor-sugar associations with paired artificial KC activation (green circle). Appetitive memory was either tested immediately after training, or 10min, 15min, 30min and 45min after training. (A^II^,B) H24>ChR2-XXL larvae trained under red light (**“sugar only“**; grey) showed significant performance scores immediately after training as well as 10min after training. No memory was detectable 15min or later after training. H24>ChR2-XXL larvae trained with blue light exposure and sugar stimulus during training (**“blue light + sugar“**; green) showed significant memory 45min after training. (C) Schematic of the mushroom body circuit focusing on the connectivity with dopaminergic input neurons. Recurrent signaling between Kenyon cells and dopaminergic pPAM neurons exists enabling ongoing activity of the dopaminergic neurons and facilitation of KC-to-MBON (KC: Kenyon cell, MBON: mushroom body output neuron) signaling. ChR2-XXL: channelrhodopsin2-XXL; ca: calyx; KCs: Kenyon cells; la: lateral appendix; MBON: mushroom body output neuron; ml: medial lobe; OCT: octanol; ped: peduncle; pn: projection neuron; vl: vertical lobe. Significance levels: p>0.05 n.s., p<0.05 *, p<0.01 **, p<0.001 ***.

## 4. Discussion

The MBs are multimodal integration centers. Beside their main function in olfactory learning and memory, MBs are involved in the control of locomotor behavior, sleep, visual learning, courtship conditioning, as well as context-dependent associations^42^. Also in the *Drosophila* larva, several studies accurately demonstrated a role of MB KCs for olfactory learning and memory^5^: classical genes involved in learning and memory like *rutabaga* and *dunce*, both part of the cAMP signaling pathway, are expressed in larval KCs^43,44^. Mutants for both genes are impaired in olfactory learning and memory. Moreover, input into the MBs is required to establish a memory during training, while output of MB KCs is necessary for memory retrieval^3,44^. These and other findings allowed to postulate a minimal and functional MB network model. However, the recent larval MB connectome described several novel signaling routes besides the classical MBIN-to-KC-to-MBON circuit^11^, which is in line with new data for the adult MB circuit^20,41,45–47^. Especially the role of MB KCs has been underestimated, as they form KC-to-KC synapses and provide a recurrent loop with MBINs. As the function of this recurrent loop is not yet known in larvae, we explored the function of this new circuit motif for its role in larval olfactory learning and memory in this study.

Using two different MB-specific driver lines we found that optogenetic KC activation is sufficient to induce olfactory appetitive memory. The artificial activation of all KCs did not alter odor processing, sugar processing, or larval locomotion. Based on these findings, we suggest to extend the current appetitive olfactory learning circuitry in the following way: during larval odor-sugar conditioning, CS (odor) information is mediated by projection neurons to the MB calyx, while simultaneously US (sugar) information is mediated via dopaminergic pPAM neurons to the medial lobe. The coincident odor-dependent Ca^2+^ influx and DA release induces cellular plasticity (in terms of short-lasting memories; this study) or synaptic plasticity between KCs and MBONs (for longer-lasting memories)^4^. Our new data suggest, that in addition pPAM neurons receive input from KCs during training. Optogenetic activation of KCs was sufficient to activate the dopaminergic pPAM neurons indicating a functional connection of KCs-to-pPAM, which was anatomically identified in the larval MB connectome^11^. However, whether the KC-to-pPAM signaling is direct or indirect remains still elusive. Yet, during odor-sugar training a recurrent circuit of KCs and pPAMs could become active, instruct and consolidate an appetitive olfactory memory. Indeed, when this loop was blocked, either through DopR1 knockdown in KCs or ablation of pPAM neurons, the induction of appetitive memory by optogenetic activation of KCs was impaired.

In adult *Drosophila*, activity in dorsal paired median neurons is necessary during the consolidation phase after training to form a persistent memory^48^. However, also during training ongoing activity and feedback signaling is necessary for olfactory learning and memory as impaired cholinergic input from KCs to dopaminergic MBINs reduces odor-electric shock memories in adult *Drosophila*^41^. In this sense, KC-to-DAN feedback signaling seems to affect neuronal activity in MBINs and by that may enhance the release of DA to the MBs beyond the level induced by the US stimulus alone^41^. Likewise, dopaminergic MBINs receive excitatory input from MBONs during courtship training and training for odor-sugar associations, which in turn leads to a persistent activity of these DANs and an increase in DA release onto the MBs. Thus, MBONs are not only necessary for memory retrieval, but also for memory acquisition and consolidation. Consequently, a longer-lasting memory is formed by the facilitation of KC-to-MBON signaling through the recurrent MBON-to-DAN feedback loop^20,47^. Taken together, within the MB circuitry direct recurrent KC-to-DAN loops together with one step recurrent KC-MBON-DAN loops exist to shift an initial nascent memory into a stable memory.

### Do recurrent circuits within the larval mushroom bodies allow the formation of different types of memories?

Our results indicate that a paired exposure to sugar and optogenetic activation of KCs increases the persistence of odor-sugar memory. Larvae trained for only odor-sugar associations under red light showed memory expression up to 10min. 15min after training the appetitive olfactory memory was abolished. On the contrary, artificial activation of KCs combined with sugar exposure during training was sufficient to induce a longer-lasting appetitive olfactory memory. Memory expression was still detectable 45min after training. Interestingly, in adult *Drosophila* activity in dopaminergic MP1 neurons is necessary in the first 30-45min after training to trigger long-term memory formation^21,49^. Thus, we assume that an increased and persistent activity within the MB network, including the KC-to-pPAM recurrent circuit signaling, may likewise support the formation of longer-lasting memories in the *Drosophila* larva. It is tempting to speculate that the activity level in dopaminergic pPAM neurons, which is increased via the KC-to-pPAM signaling through optogenetic activation of KCs in our study, is crucial to drive the conditional behavioral output. Different activity levels set a specific level of intracellular cAMP, either triggering a short-lasting memory (by a cAMP-dependent transient increase in protein kinase A (PKA)) or longer-lasting memories (by more stable cAMP-dependent elevation of PKA)^50,51^. Changes in DA neuronal activity (presumably controlled by the existing feedback loops) may provide a steady update about the internal state and the expected gain from memory expression based on available environmental stimuli. The type of internal information may thus be appropriate to regulate the type of memory which is formed. In short, under food restriction adult flies preferentially show anesthesia-resistant memory rather than long-term memory, as this consolidated memory form is independent of energy-demanding protein biosynthesis and therefore at low cost^52,53^. Further, in both flies and larvae, neurons expressing the neuropeptide dNPF (*Drosophila* neuropeptide F; an orthologue of mammalian NPY) modulate memory processes to match information about the current reward stimulus with the internal state of the animal. A conditional activation of dNPF neurons during training reduced the acquisition of odor-sugar memories in the *Drosophila* larva^54^. Odor-sugar learning in flies is most efficient when flies are food deprived. dNPF neurons modulate the expression of odor-sugar memories via feedforward inhibition through dopaminergic MBINs and the MBON circuitry. In hungry flies, specific MBONs (MV2) are inhibited by the action of dNPF neurons through MP1 DANs, which elicits the expression of odor-sugar memories and thus conditional approach behavior^55^.

Thus, DANs innervating different compartments of the MBs may provide a dynamic intrinsic and extrinsic neuromodulatory system: depending on experience and the internal state, the same sensory input (e.g. a sugar stimulus) may drive a different conditional behavioral output (either approach, avoidance or indifference) and different temporal forms of associative olfactory memories (short-term memory, anesthesia-resistant memory or long-term memory)^45,56^.

Our assumption is now that a recurrent signaling loop within the larval MB circuit allows the larva to constantly adapt appetitive memory formation and retrieval to the current internal state and external situation, mainly based on the modulation of ongoing activity in pPAMs, and by that to incorporate whether the expected gain is sufficient to store and recall a long-lasting associative memory. In humans, DA was shown to modulate a post-encoding consolidation process with respect to episodic memory persistence^57^. The molecular consolidation process is based on DA-dependent protein-synthesis, which elicits long-term plasticity in the hippocampus. Similarly, in rodents the activation of different DA receptors in the hippocampus is critical at or around the time of memory encoding to modulate memory persistence^58^. Thus, activation of recurrent signaling routes within a neuronal memory circuit via DANs and the resulting sustained activity in these neurons appears as a conserved mechanism to consolidate memories in insects and vertebrates.

## Supplement figures

**Supplement figures:**
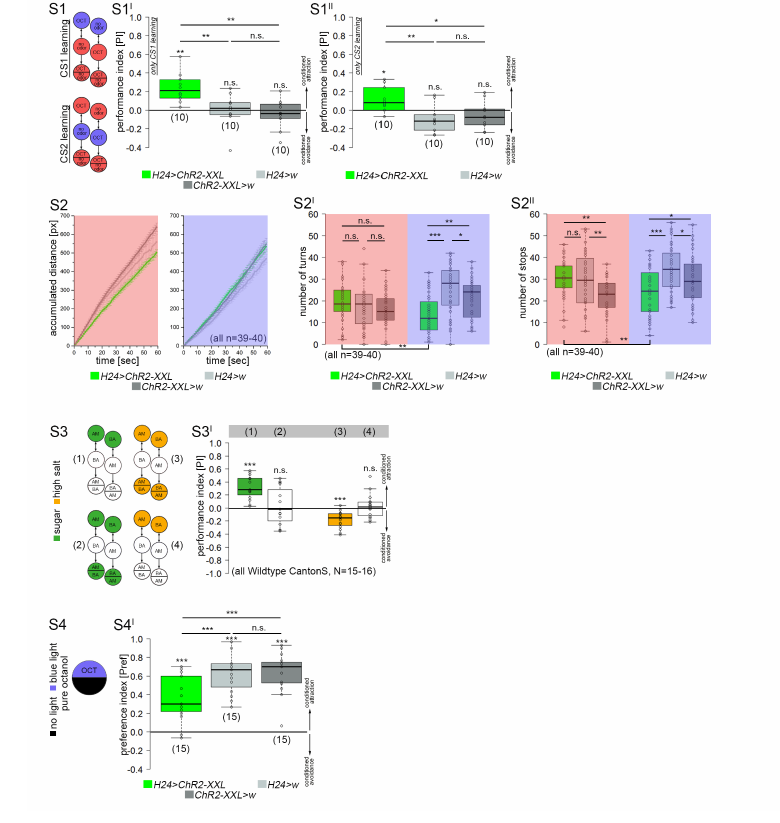
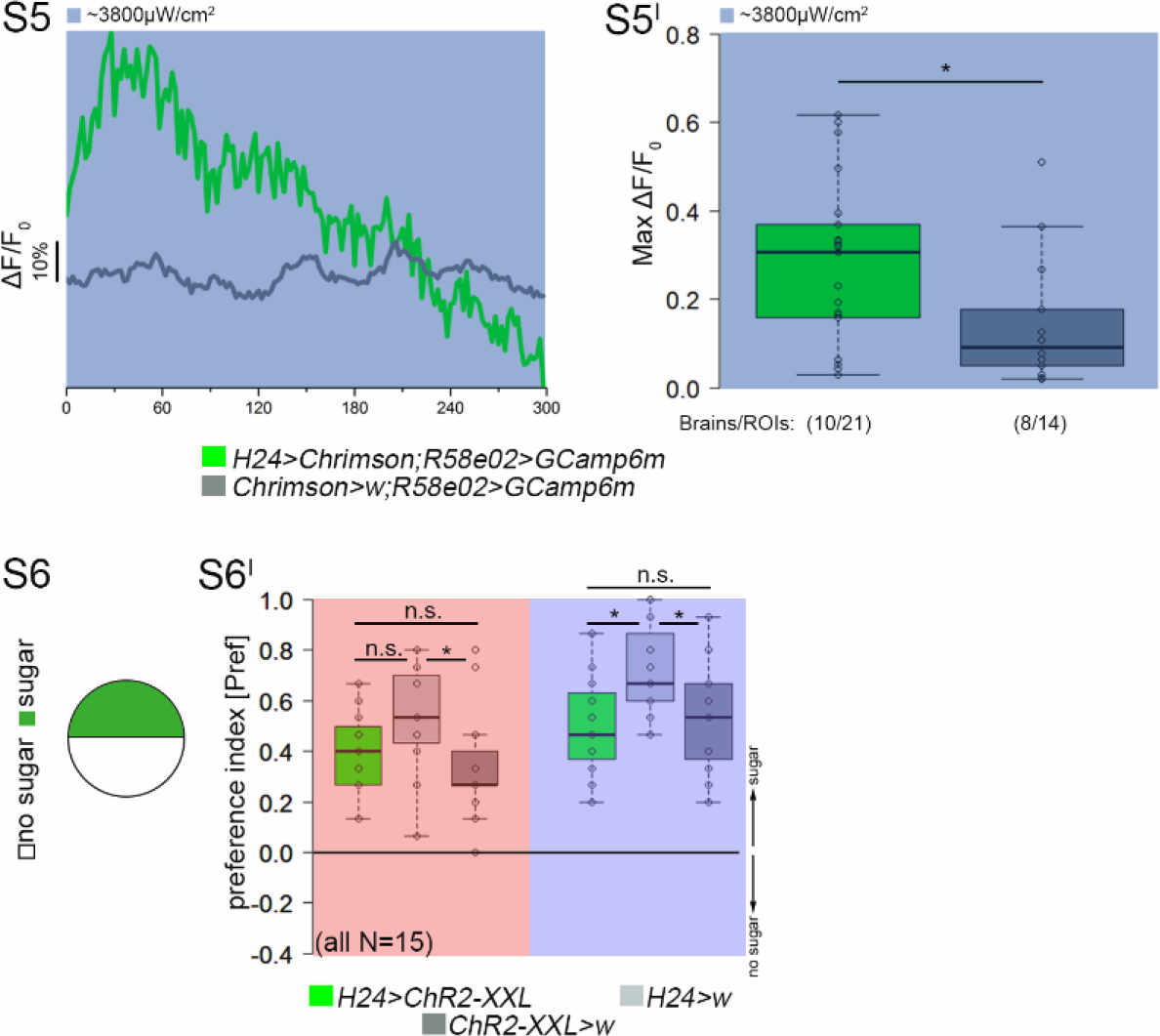
(S1-S1^II^) Performance scores were independent of whether the first odor (CS1+; OCT or “no odor“) or the second odor (CS2+; OCT or “no odor“) was coupled to optogenetic activation of KCs during KC-substitution learning. (S2) General locomotion was unaffected by optogenetic activation of all KCs as H24>ChR2-XXL larvae crawled normal distances over time compared to genetic controls. Experimental larvae showed a reduced number of turns (S2^I^) and stops (S2^II^) due to blue light exposure. (S3-S3^I^) Wildtype larvae express appetitive olfactory memories not in presence of the appetitive US (e.g. sugar (green)) in the test situation, while aversive memory expression is only present when the aversive US (e.g. high salt (orange)) is absent in the test situation. (S4-S4^I^) H24>ChR2-XXL larvae show a reduced light avoidance compared to genetic controls when they are challenged to decide between their innate OCT and darkness preference versus an optogenetically induced internal reward signaling. (S5) pPAM neurons respond with a significant increase in intracellular Ca^2+^ levels (S5C^I^) to an optogenetic activation of MB KCs using UAS-Chrimson (green; illumination with blue light (475nm) was already sufficient for optogenetic activation via Chrimson) verifying a functional KC-to-DAN feedback connection. An obvious increase in Ca^2+^ levels was absent in genetic controls (grey). (S6-S6^I^) H24>ChR2-XXL larvae showed normal innate preference for fructose compared to genetic controls. AM: amylacetate; BA: benzaldehyde; ChR2-XXL: channelrhodopsin-XXL; CS: conditioned stimulus; OCT:octanol. Significance levels: p>0.05 n.s., p<0.05 *, p<0.01 **, p<0.001 ***.

## 6. Material and Methods

### 6.1 Fly strains

Flies were cultured according to standard methods. In short, vials were kept under constant conditions with 25°C and 60% humidity in a 12:12 light:dark cycle. Driver lines used in this study were *H24-Gal4* (chromosome II), *Ok107-Gal4* (IV), *R58E02-Gal4* (III), and *R58E02-lexA* (II). UAS lines included in this study were 10xUAS-*IVS-myr::GFP* (II), UAS-*mCD8::GFP* (II), UAS-*ChR2-XXL* (II), 20xUAS-*CsChrimson* (II), UAS-*DopR1-RNAi* (III), *lexAop-reaper* (II), and 13x*LexAop*-*IVS-GCamp6m*/20xUAS-*CsChrimson* (II,III). Genetic controls were obtained by crossing Gal4-driver/Lex-driver and UAS-effector/LexAop-effector lines to *w^1118^.*

### 6.2 Behavioral experiments

#### 6.2.1 Associative conditioning

Appetitive olfactory learning was tested using standardized, previously described assays^5,59^. Learning experiments were performed on plates filled by a thin layer of pure 1.5% agarose solution (Roth, 9012-36-6). We mainly used a one-odor reciprocal training design^22^, where larvae were exposed to 10μl of 1-octanol (OCT, Sigma, 111-87-5). We also performed one experiment using the two-odor reciprocal training design, where 10μl of amylacetate (AM, Merck, 628-63-7) was exposed to the larvae opposing OCT. Odorants were loaded into custom-made Teflon containers (4.5mm diameter) with perforated lids.

For KC-substitution learning experiments, a first group of 30 larvae was trained to associate OCT to blue light exposure (OCT+). After 5min the larvae were transferred to a second Petri dish containing no odor under red light (NO). Simultaneously, a second group of larvae was trained reciprocally with blue light exposure coupled to no odor (NO+/OCT). After three training cycles, larvae were immediately transferred to the test plate, where OCT was presented on one side of the dish. To test the persistence of the established memory, larvae were tested 10min, 15min, 30min, or 45min after the training. After 3min, larvae were counted in the OCT side (#OCT), the no odor side (#NO), and in the neutral zone (for further details, a video is provided in^59^). The procedure was similar for the two-odor reciprocal training design (OCT+/AM and OCT/AM+). The preference index was calculated by subtracting the number of larvae on the OCT side from the number of larvae on the no odor side (or AM side), divided by the total number of animals (including the larvae counted in the neutral zone).

a. PREFOCT+/NO = (#OCT − #NO) / #TOTAL
b. PREFOCT/NO+ = (#OCT − #NO) / #TOTAL
c. PI = (PREFOCT+/NO − PREFOCT/NO+) / 2

Subsequently, we calculated the performance index (PI). Positive PIs indicate appetitive learning, while negative PIs indicate aversive learning.

For optogenetic manipulation in all behavioral experiments (using UAS-*ChR2-XXL*) we used 475nm light-emitting diodes (LED) with a light intensity of ˜ 1300µW/cm^2^, or red light with 620nm and the intensity of 50 µW/cm^2^ for UAS-*Chrimson*. To induce optogenetic activation, the light-emitting diodes were placed ˜45cm above the Petri dish, while all other steps of the experimental procedure were done under red light (or complete darkness for UAS-*Chrimson*).

#### 6.2.2 Olfactory preference tests

To test larvae for their innate odor response, an odor container was placed on one side of the Petri dish containing 1.5% agarose to induce a choice test. For optogenetic manipulation the Petri dish was placed in blue light, for control experiments the choice test was done under red light. The preference index was calculated by subtracting the number of larvae on the odor side (#Odor) from the number of larvae on the no odor side (#NO), divided by the total number of animals (including the larvae counted in the neutral zone). In each test, we used a group of 30 larvae.

PREFOdor/NO = (#Odor − #NO) / #TOTAL

Negative PREF values indicate an avoidance of the odor, whereas positive PREF values represent an attractive response.

#### 6.2.3 Gustatory preference tests

To test larvae for their innate gustatory response during optogenetic activation, one half of the Petri dish was filled with 1.5% pure agarose, while the other half was filled with 1.5% agarose containing 2M fructose (Roth, 57-48-7). For optogenetic manipulation the Petri dish was placed in blue light, for control experiments the choice test was done under red light. The preference index was calculated by subtracting the number of larvae on the sugar side (#Sugar) from the number of larvae on the no sugar side (#NS), divided by the total number of animals (including the larvae counted in the neutral zone). In each test, we used a group of 30 larvae.

PREFSugar/NS = (#Sugar − #NO) / #TOTAL

Negative PREF values indicate sugar avoidance, whereas positive PREF values represent an attractive response.

#### 6.2.4 Light preference tests

To test larvae for their response to blue or red light, respectively, (and by that to optogenetic manipulation), the Petri dish containing 1.5% agarose was covered by a lid, divided in two transparent and two shaded quarters, respectively. The preference index was calculated by subtracting the number of larvae on the dark side (#DS) from the number of larvae on the illuminated side (#Light), divided by the total number of animals (including the larvae counted in the neutral zone). In each test, we used a group of 30 larvae.

PREFDS = (#DS − #Light) / #TOTAL

Positive PREF values indicate light avoidance, whereas negative PREF values represent approach towards the illuminated side.

### 6.3 Locomotion assay

For the locomotion assay we used the FIM (FTIR-based Imaging Method) tracking system as described in^27^. Recordings were made by a monochrome industrial camera (DMK27BUP031) with a Pentax C2514-M objective in combination with a Schneider infrared pass filter, and the IC capture software (www.theimagingsource.com). To analyze larval locomotion, a group of 10 larvae was recorded on 1.5% agarose for two minutes. During the first minute, larvae were exposed to red light, while they were exposed to blue light for the second minute for optogenetic activation of KCs. We analyzed the following parameters using the FIM track software: accumulated distance, velocity, number of stops, and number of bendings.

### 6.4 Immunofluorescence

Immunofluorescence studies were performed as described in^60^. In short, 5-6 day old larvae were dissected in phosphate buffer saline (PBS) or HL3.1 (pH 7.2)^61^, fixated in 4% paraformaldehyde in PBS for 40min, washed four times in PBS with 0.3% Triton-X 100 (PBT), and afterwards blocked in 5% normal goat serum in PBT. Specimens were incubated in primary antibody solution containing polyclonal rabbit anti-GFP antibody (A6455, Molecular Probes, dilution 1:1000) in blocking solution for one night at 4°C. Then brains were washed six times in PBT and incubated for one night at 4°C in secondary antibody solution containing goat anti-rabbit Alexa 488 (Molecular Probes, dilution 1:250). Finally, specimens were rinsed six times in PBT and mounted in 80% glycerol in PBS. Until scanning with a Leica SP8 confocal light scanning microscope, brains were stored in darkness at 4°C. Image processing was performed with Fiji^62^ and Adobe Photoshop CS6 (Adobe Systems, San Jose, CA).

### 6.5 Functional imaging

To monitor intracellular Ca^2+^ levels in dopaminergic neurons of the pPAM cluster in response to the optogenetic activation of MB KCs, LexAop-*GCaMP6m* was expressed via *R58E02-LexA* and UAS-*ChR2-XXL* via *H24-Gal4*. 1-2 larval brains were dissected and subsequently put in a Petri dish containing 405μl hemolymph-like HL3.1 saline solution. Before the experiment, brains maintained for around 30min for settling. Specimens were imaged with an AXIO Examiner D1 upright microscope (Carl Zeiss AG, Germany) with a Cairn Twin-Cam LS image splitter, two pco.edge 4.2 sCMOS cameras, and a SPECTRA-4 light engine. Images were taken with a Zeiss W Plan-Apochromat x20/1.0 or x40/1.0 water immersion objective. During monitoring, brains were first excited with 475nm light and an exposure time of 60ms at 4x binning with an intensity of around 700μW/cm^2^. After 5min, light intensity was increased to around 3800μW/cm^2^ for optogenetic activation of MB KCs. We continued the monitoring of Ca^2+^ levels in pPAM neurons for 5min. Similarly, MB KCs were optogenetically activated via UAS-*Chrimson*. Here, we recorded Ca^2+^ levels in *R58E02>GCaMP6m* brain with 475nm light and an intensity of 3800μW/cm^2^, but without pre-exposure of a lower light intensity.

### 6.6 Statistical methods

Data was analyzed for normal distribution using the Shapiro-Wilk Normality test. To test against chance level, a t-test was used for normally distributed data, a Wilcoxon Signed Rank test for not normally distributed data. For the comparison between genotypes, a pairwise t-test was used for normally distributed data, a pairwise Wilcoxon Rank Sum test was used for not normally distributed data. Pairwise tests included the Bonferroni-Holm correction. All statistical analyses were done with R Studio version 0.99.896 (www.r-project.org). Data plots were done with OriginPro 2016G, b9.3.226 (www.originlab.com). Data is mainly presented as box plots, with 50% of the values of a given genotype being located within the box, and whiskers represent the entire set of data. No data was excluded. Outliers are indicated as open circles. The median performance index is indicated as a thick line within the box plot. For the persistence of memory, data is also presented as a line chart, with mean values and the standard error of the mean. Significance levels between genotypes shown in the figures refer to the raw p-values obtained in the statistical tests.

## Acknowledgements

We thank Konrad Öchsner, for technical assistance, Wolf Hütteroth, Katharina Eichler, Björn Brembs, Thomas Raabe, Tim Humberg, and Martin Strube-Bloss for fruitful discussions and/or comments on the manuscript. We thank Robert Kittel, Georg Nagel, Katharina Eichler, Vivek Jayaraman, the Vienna *Drosophila* resource center, and the Bloomington Stock center for providing flies. The authors declare no competing interests. R.L., A.S.T. and D.P. conceived and designed the experiments. R.L., M.S., M.P., J.H., D.S. and D.P. performed the experiments. R.L., M.S., C.W. and D.P. analyzed the results. R.L., C.W., A.S.T., and D.P. wrote the manuscript. This work was supported by the PostDoc Plus fellowship (to D.P.) and a PhD fellowship (to R.L.) from the German Excellence Initiative to the Graduate School of Life Sciences, University of Würzburg, and by the Deutsche Forschungsgemeinschaft (TH1584/1-3 to A.S.T., INST 93/824-1 LAGG to C.W., and DP1979/2-1 to D.P.) and intramural funds by the University of Würzburg (to C.W.).

